# Curcumin delays cytokinesis in fission yeast by targeting the septin ring

**DOI:** 10.1101/2025.06.02.657428

**Authors:** Varmila Kulasegaram, Daphne Alves Dias, Anna Okorokova-Façanha, Qian Chen

## Abstract

Curcumin is the active compound of one of the most widely used spices in the world. It has also been highly valued as a traditional health supplement in many South Asian countries for millennia. More recently, this plant extracted small molecule has attracted strong attention for its therapeutic potential. Nevertheless, the molecular and cellular targets of curcumin remain unknown. Here, we undertook a novel imaging-based approach to determine the intracellular distribution of curcumin using the model organism fission yeast *Schizosaccharomyces pombe* by taking advantage of the intrinsic fluorescence of curcumin. Live fluorescence microscopy revealed for the first time that curcumin, at a concentration of just one micromolar, formed a narrow circumferential ring around the equatorial plane of dividing cells within minutes after being added to the yeast culture. The intensity of this ring increased proportionally to the concentration of curcumin and gradually over time. The curcumin ring co-localized both spatially and temporally with septin ring and the exocyst complex at the equatorial plane during cytokinesis. Deletion of one of the septin genes *spn4* reduced the frequency of curcumin ring by 74%. Micromolar concentrations of curcumin slowed down the contractile ring constriction by up to 49% in a dosage dependent manner. Besides fission yeast, curcumin similarly targets the division plane of two other yeasts, *Saccharomyces cerevisiae* and *Candida albicans*. Thus, curcumin, originated from plants, targets the septin cytoskeleton to delay yeast cytokinesis, suggesting its potential as both an antifungal therapeutic option and a fluorescence probe for yeast septins.

## Introduction

Curcumin is the main active compound of a widely used spice around the world. This spice, also called turmeric, is isolated from the root of the ginger family plant *Curcuma longa*. Originated in India, this yellow spice has been used in South Asia for millennia. Over the last few centuries, curcumin has gained even more popularity, adopted by many other countries around the world.

Besides being used in food, curcumin has a long history of being used as either dye, or herbal medicine, or cosmetics. More recently, curcumin has especially attracted strong interest in biomedical research for its therapeutic potential as an anti-tumor, anti-oxidant or anti-inflammatory supplement (Gupta et al., 2013; Stepien et al., 2020; Zhou et al., 2011). The heightened interest in curcumin can be gleaned from the large number of scientific studies related to this small molecule. For example, in 2024 alone, more than 2,400 papers, related to curcumin, appeared in the publicly available database PubMed. Despite such strong interest in the potential of curcumin for human health, the cellular and molecular targets of this small molecule remain obscure.

Yeasts have been instrumental in elucidating the cellular mechanisms of various bioactive compounds. This is due to that many of their signaling pathways are conserved in animal cells and there are powerful genetic tools available in yeasts (Botstein and Fink, 2011). In particular, the budding yeast *Saccharomyces cerevisiae* has helped identify some of the potential molecular targets of curcumin. In the budding yeast cells treated with high (100 µM) concentrations of curcumin, this small molecule accumulates in the endoplasmic reticulum (ER) (Minear et al., 2011). Curcumin chelates intracellular iron in budding yeast, potentially disrupting the iron homeostasis (Azad et al., 2014; Minear et al., 2011). In addition, several reports also documented the pleiotropic effects of curcumin in *S. cerevisiae*. For example, curcumin can activate the MAPK kinase cascade (Azad et al., 2014). It may also delay cell cycle progression (Minear et al., 2011). Finally, curcumin can extend the chronological lifespan of yeast (Stepien et al., 2020). Overall, it remains unclear what the main cellular and molecular targets of curcumin are in yeast.

Fission yeast *Schizosaccharomyces pombe* is another yeast that has served as an excellent eukaryotic model organism (Hoffman et al., 2015). Despite some similarities, *S. pombe* and *S. cerevisiae* differ significantly from each other in many aspects. This is not surprising, considering that, they diverged from each other more than 300 million years ago (Rhind et al., 2011). In growth, the cylindrical-shaped fission yeast cells grow through tip extension, unlike the spherical-shaped budding yeast cells. In terms of the cell cycle, the *S. pombe* cells divide through fission, instead of budding. In cytokinesis, fission yeast cells assemble an essential actomyosin contractile ring at the equatorial division plane, followed by the cleavage furrow ingression and the septation. Unlike *S. cerevisiae*, dividing fission yeast cells split into two equal sized daughter cells. To our knowledge, few have studied the cellular effects of curcumin in fission yeast.

Here, we employed a novel imaging-based approach to trace the intracellular distribution of curcumin in fission yeast cells. To our surprise, we discovered that curcumin marks a circumferential ring at the equatorial plane even at a concentration of just one micromolar. This curcumin ring co-localized with the septin cytoskeletal structure; septin ring during cytokinesis. Deletion of one of the septin genes inhibited the formation of the curcumin ring. Even low concentrations of curcumin inhibited the progression of cytokinesis in a dosage-dependent manner. Finally, we found that curcumin marks the equatorial division plane of two other yeasts, *S. cerevisiae* and *Candida albicans*. The discovery of the unique septin-dependent curcumin ring in yeast cells and its disruptive effect against yeast cytokinesis demonstrate both the therapeutic potential of curcumin and the promising future of using curcumin as a fluorescent probe to study the yeast septin cytoskeletal structures.

## Results and discussion

One unique property of curcumin is its intrinsic fluorescence, which allows direct visualization of this small molecule. Curcumin (C21H20O6; diferuloylmethane) has a molecular weight of 368 g/mol (Fig. 1A). It can be excited at 410-500nm with the emission peaking at 500-550nm, depending on the solvent (Ferreira et al., 2023; Priyadarsini, 2009; Zsila et al., 2003). We found that curcumin dissolved easily in the fission yeast media YE5s. As expected, the 100µM curcumin supplemented media appeared as yellow under visible light (Fig. 1A). In comparison, a much lower concentration (8µM) of curcumin did not alter either the color or the transparency of YE5s (Fig. 1A). We first tested whether we could trace the intracellular distribution of this small molecule through fluorescence microscopy. We inoculated exponentially growing fission yeast cells in liquid YE5s supplemented with 0.5, 1, 2, 4 and 8 µM of curcumin respectively for 10 minutes (Fig. 1B). Then we imaged these cells using a confocal microscope, with the excitation/emission wavelength of the laser set at 488/525 nm. Little or no fluorescence was detected in the cells treated with either the vehicle (DMSO) or 0.5µM curcumin (Fig. 1B). In comparison, at 1µM, a bright fluorescent band appeared at the equatorial plane of the yeast cells. Except for this band, no other intracellular structures appeared as fluorescent (Fig. 1B). The fluorescence intensity of these bands increased, correlated with the increasing concentrations of curcumin (Fig. 1B). At 8µM, in addition to the equatorial band, the cytoplasm of some cells also became fluorescent (Fig. 1B). At these relatively low concentrations that we used, curcumin had no observable effects on the morphology of fission yeast cells (Fig. 1B). Continued incubation of the yeast cells in curcumin, up to 70 mins, further increased the fluorescence intensity of the curcumin band (Fig. 1C). When reconstructed from the Z-stacks for a head-on view, the band emerged as a ring encircling the equatorial plane (Fig. 1C), which we called “curcumin ring”. To determine whether this curcumin ring specifically targets the dividing cells, we prevented the cells from entering mitosis by inactivating the phosphatase Cdc25 that is essential for the G2 to M transition, using a temperature-sensitive *(cdc25-22)* mutant (Gould et al., 1990). At the restrictive temperature of 36°C, these mutant cells failed to divide, thus becoming much longer than the wild-type cells (Fig. 1D). No curcumin ring was detected in these mutant cells (Fig. 1D). Interestingly, the intensity of the curcumin ring in the wild type cells also weakened significantly at 36°C (Fig. 1D), suggesting that the curcumin ring is likely temperature dependent. We concluded that low concentrations of curcumin form a ring circumscribing the equatorial plane of dividing fission yeast cells.

**Figure 1.**
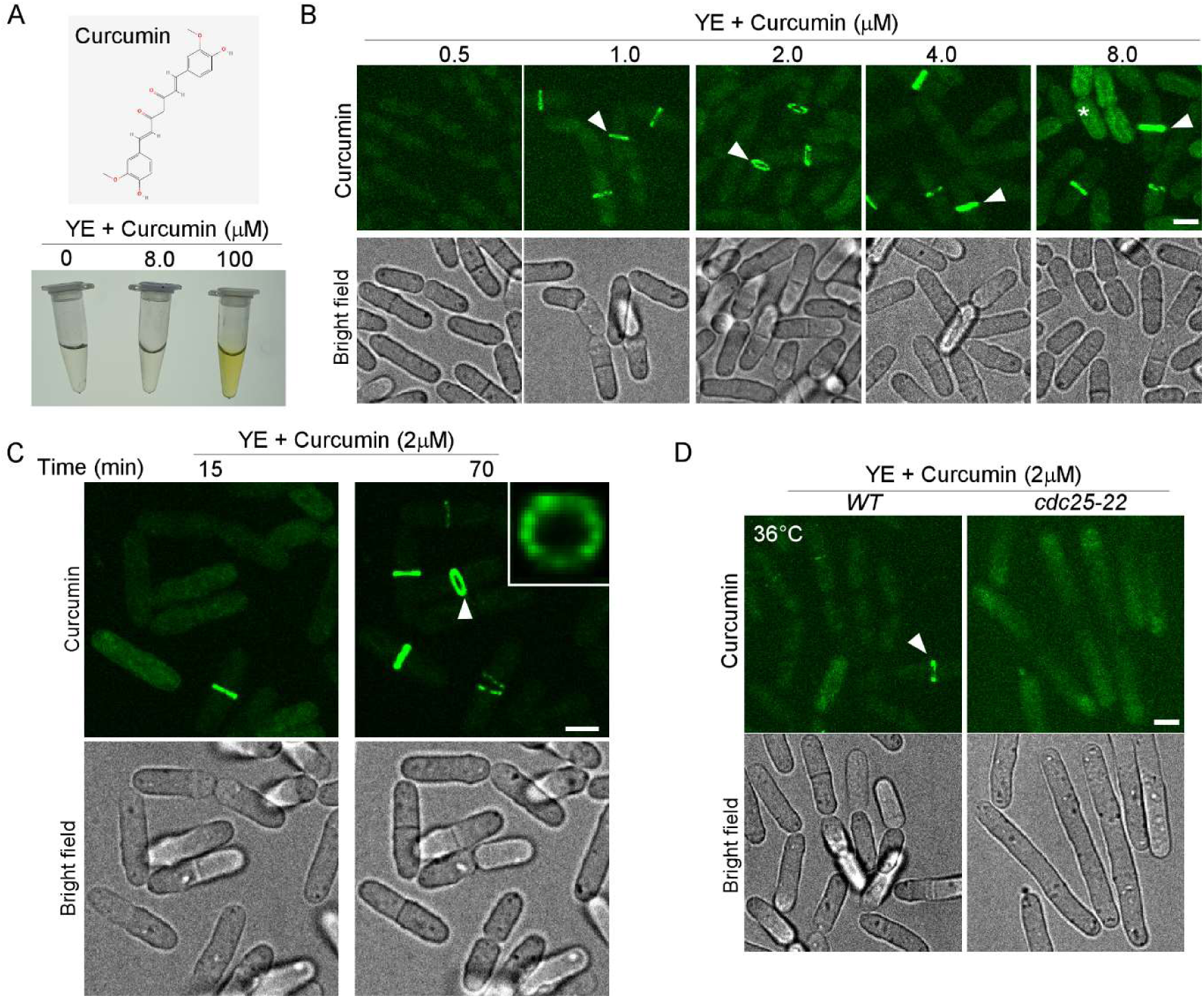
Curcumin targets a ring-like structure at the equatorial plane of dividing fission yeast cells. (A) Top: Chemical structure of curcumin (from PubChem). Bottom: Photos of YE5s media in microcentrifuge tubes, supplemented with 0, 8 and 100µM curcumin respectively. At 100 µM, the curcumin supplemented media appeared as yellow under light. (B) Micrographs of fission yeast cells in the presence of various concentrations of curcumin after 10 min incubation. Asterisk: a cell with cytosolic fluorescence from curcumin. (C) Micrographs of fission yeast cells in the presence of 2µM curcumin for 15 and 70 mins respectively. Insert: 3D reconstructed head-on view of the curcumin band (arrowhead). (D) Micrographs of the wild type (left) and *cdc25-22* mutant (right) cells in the presence of curcumin at the restrictive temperature of 36°C. Arrowhead: curcumin band. Representative micrographs from three independent biological repeats are shown. Bar: 5 µM.

At the division plane, curcumin can potentially target many components of the yeast cytokinetic machinery to form the ring. To determine which one of them is the likely target of curcumin, we co-localized the curcumin ring with the septum, the plasma membrane, and the cytoskeletal structures at the equatorial division plane respectively (Fig. 2). First, we started with the septum, the newly synthesized cell wall at the division plane, based on our assumption that curcumin has to cross this barrier before entering the cells. Live microscopy showed the curcumin ring was present in many cells whose septum, stained by the fluorescence dye calcofluor (Ribas and Cortés, 2016), had not yet been assembled (Fig. 2A). In those presumed late cytokinesis cells, the septum appeared as a disc separating daughter cells, observed from the cross-section view of the division plane. This was in contrast to the ring formed by curcumin in these cells (Fig. 2A, magnified head-on views). To confirm that the septum is spatially distinct from the curcumin ring, we examined the co-localization between curcumin and the glucan synthase Bgs1, which synthesizes the 1,3-β-glucan at the septum (Cortes et al., 2002), tagged with tdTomato. Bgs1-tdTomato did appeared as a ring in many dividing cells, overlapping with the curcumin ring (Fig. 2B). However, in many other dividing cells, Bgs1-tdTomato appeared as a disc separating the daughters, just like the septum but distinct from the curcumin ring (Fig. 2B, magnified head-on views). Therefore, we concluded that the septa is unlikely to be a target of curcumin. The plasma membrane at the division plane was the next potential target that we examined. We used a putative ion channel Trp663 tagged with mCherry to label the plasma membrane at the equatorial division plane (Malla et al., 2022) (Fig. 2C). In most cells expressing Trp663-mCherry, this fluorescent protein also formed a disc separating the daughter cells, unlike the ring marked by curcumin (Fig. 2C, magnified head-on views). Therefore, the curcumin ring does not target either the septa or the plasma membrane at the equatorial division plane.

**Figure 2.**
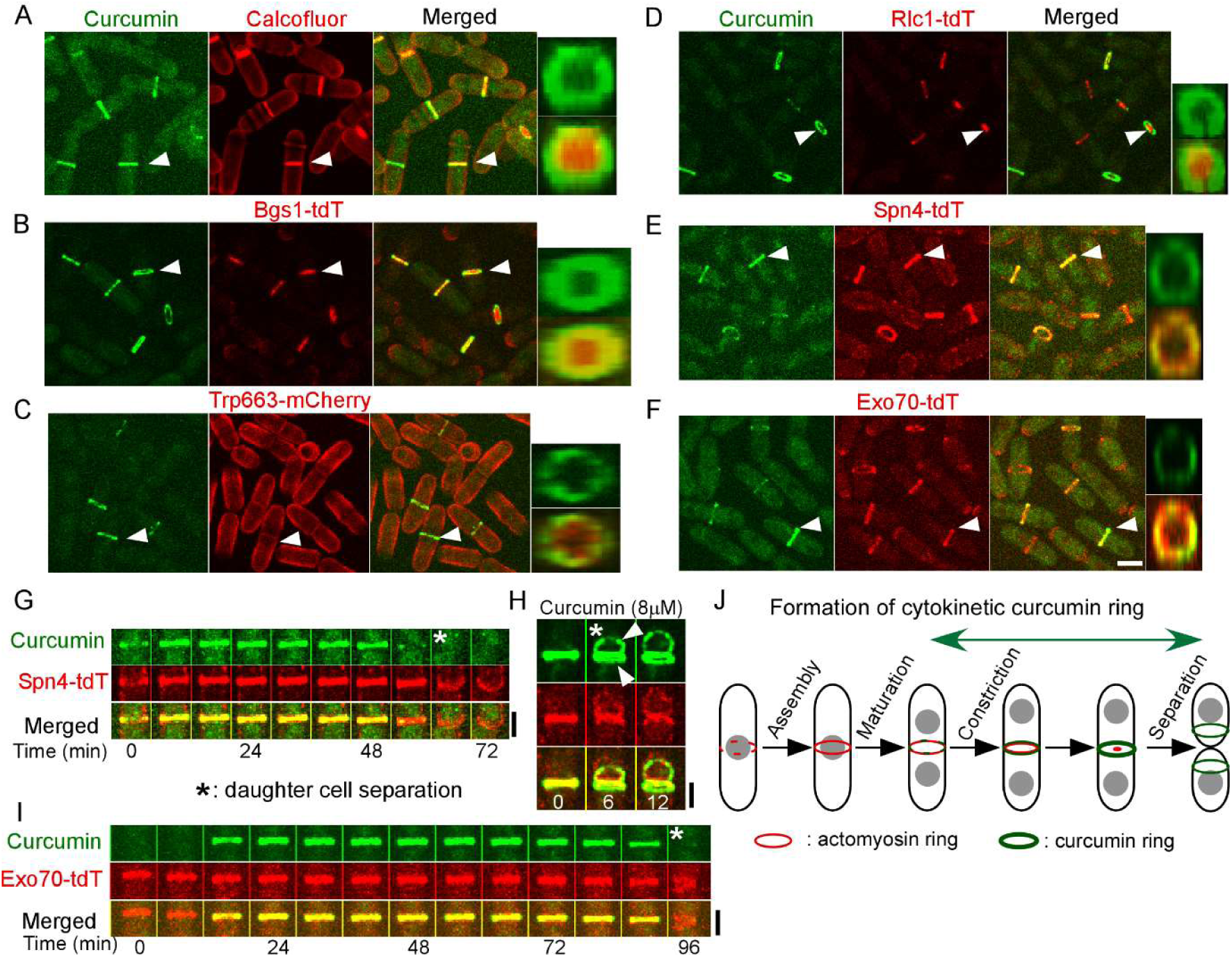
The curcumin ring co-localizes with the septin ring throughout cytokinesis. (A-F) Co-localization between curcumin (2µM, green) and various components (red) of the yeast cytokinetic machinery. Arrowhead: the curcumin ring reconstructed for magnified head-on views in the boxes (top: curcumin only, bottom: merged) on the right. (A) Micrographs of the wild type cells in the presence of calcofluor that stained the septum. (B-F) Micrographs of the cells expressing Rlc1-tdTomato (tdT, for the contractile ring), Bgs1-tdT (for glucan synthase), Trp663-mCherry (for the plasma membrane), Spn4-tdT (for the septin ring), and Exo70-tdT (for the exocyst complex) respectively. The curcumin ring co-localized with both the septin ring and the exocyst complex during cytokinesis. (G-I) Time series of the division plane of a cell expressing either Spn4-tdT (G and H) or Exo70-tdT (I) in the presence of either 2 (G) or 8 (H) μM of curcumin. Number: time in minutes. Arrowhead: two split curcumin rings after the daughters separated (asterisk). (J) A diagram depicting the formation of the curcumin ring during cytokinesis relative to the contractile ring assembly, maturation and constriction. Representative data from three independent biological repeats are shown. Bar: 5 µM.

This prompted us to determine whether curcumin recognizes the cytoskeletal structures at the division plan. Both actins and septins assemble into a ring at the division plane, making both potential targets of curcumin. For the former, we colocalized curcumin with the actomyosin contractile ring marked by Rlc1, the myosin regulatory light chain (Le Goff et al., 2000), tagged with the fluorescent protein tdTomato. In some dividing cells, the contractile ring and the curcumin ring did co-localize with each other (Fig. 2D). However, in the majority of the dividing cells, the contractile ring, due to their constriction, was smaller than the curcumin ring. This was apparent through the reconstructed cross-section view of the furrow in which the curcumin ring encircles the smaller contractile ring (Fig. 2D, magnified head-on views). Therefore, we also ruled out the possibility that the actomyosin contractile ring is the target of curcumin at the division plane. Next, we focused on the septin cytoskeleton. Fission yeast possesses seven septin proteins, four of which Spn1, 2, 3 and 4 express in the vegetative cells and oligomerize into a ring at the equatorial division plane (An et al., 2004). We used one of them Spn4, tagged with tdTomato, to mark the septin cytoskeleton in dividing cells. The septin ring co-localized perfectly with curcumin in all the dividing cells (Fig. 2E). Most tellingly, both the curcumin and the septin rings encircled the division plane (Fig. 2E, magnified head-on views). Consistent with this result, the exocyst complex labeled by Exo70-tdTomato, which physically interacts with the septins at the division plane (Singh et al., 2025), also co-localized with the curcumin ring in most of the dividing cells (Fig. 2F). Like the septin and curcumin rings, the exocyst circumscribed the division plane (Fig. 2F magnified head-on view). Overall, based on these co-localization studies, we concluded that the spatial distribution of curcumin overlaps with that of both the septin ring and the exocyst complex at the equatorial division plane, suggesting that these two are the most likely targets of curcumin.

Next, we took a step further to determine which one between the septins and the exocyst correlates temporally with the curcumin ring in dividing cells. We carried out time-lapse microscopy of the dividing cells expressing the fluorescent proteins markers for either the septins or the exocyst in the presence of curcumin. In the cells expressing Spn4-tdTomato, both the septin and the curcumin rings formed at the division plane at the same time, ∼60 mins before the cell separation (Fig. 2G), coinciding with the start of the contractile ring constriction. Both rings persisted until the end of the daughter cell separation without constricting, consistent with the previous reports (Tasto et al., 2003). Although uncommon, we observed that occasionally the septin ring split into two, one in each of the daughters, after the cell separation (Fig. 2G). Similarly, when we raised the curcumin concentration to 8µM to increase its fluorescence intensity, we also observed that the curcumin ring also split into two, overlapping with the split septin rings (Fig. 2H). In contrast to the septin ring, the exocyst complex, marked by Exo70-tdTomato, appeared ∼16 mins earlier than the curcumin ring at the division plane (Fig. 2I), coinciding with the assembly of the contractile ring. Both the exocyst complex and the curcumin ring maintained their diameters, until the daughter cell separated (Fig. 2I). Therefore, the curcumin ring forms just when the contractile ring starts to constrict and it maintains its diameter throughout the ring constriction (Fig. 2J), similar to the septin ring (Singh et al., 2025). Overall, formation of the curcumin ring closely resembles that of the septin ring both spatially and temporally in dividing yeast cells.

Having determined the spatiotemporal regulation of the curcumin localization at the division plane, we next tried to find out what is required for the formation of the curcumin ring. To answer this question, we inhibited cytokinesis with either small molecule drugs or genetic mutations. First, to determine whether the actomyosin contractile ring is required for the curcumin ring formation, we used Latrunculin A (LatA, 10 µM) that inhibits the actin filament polymerization. It quickly disassembled the Rlc1-tdTomato marked contractile ring into dozens of puncta at the equatorial division plane (Fig. 3A). In these cells, curcumin remained at the equatorial division plane as puncta, but there was no curcumin ring (Fig. 3A). Similarly, we used blebbistatin (20 µM), a small molecule inhibitor of the motor protein Myo2. It prevented the assembly and constriction of the contractile ring (Fig. 3A). In the dividing cells, the contractile ring appeared as strands at the division plane. Nevetheless, curcumin remained at the division plane, but it failed to form a ring (Fig. 3A). Consistent with the results of this pharmacological approach, in *myo2-E1* cells, a temperature-sensitive mutant of *myo2,* curcumin remained at the division plane as cytoplasmic puncta at 36°C (Fig. 3B), but it formed thick strands instead of rings (Fig. 3B). Therefore, we concluded that curcumin requires the actomyosin cytoskeleton to form a ring but it can target the division plane without the contractile ring.

**Figure 3.**
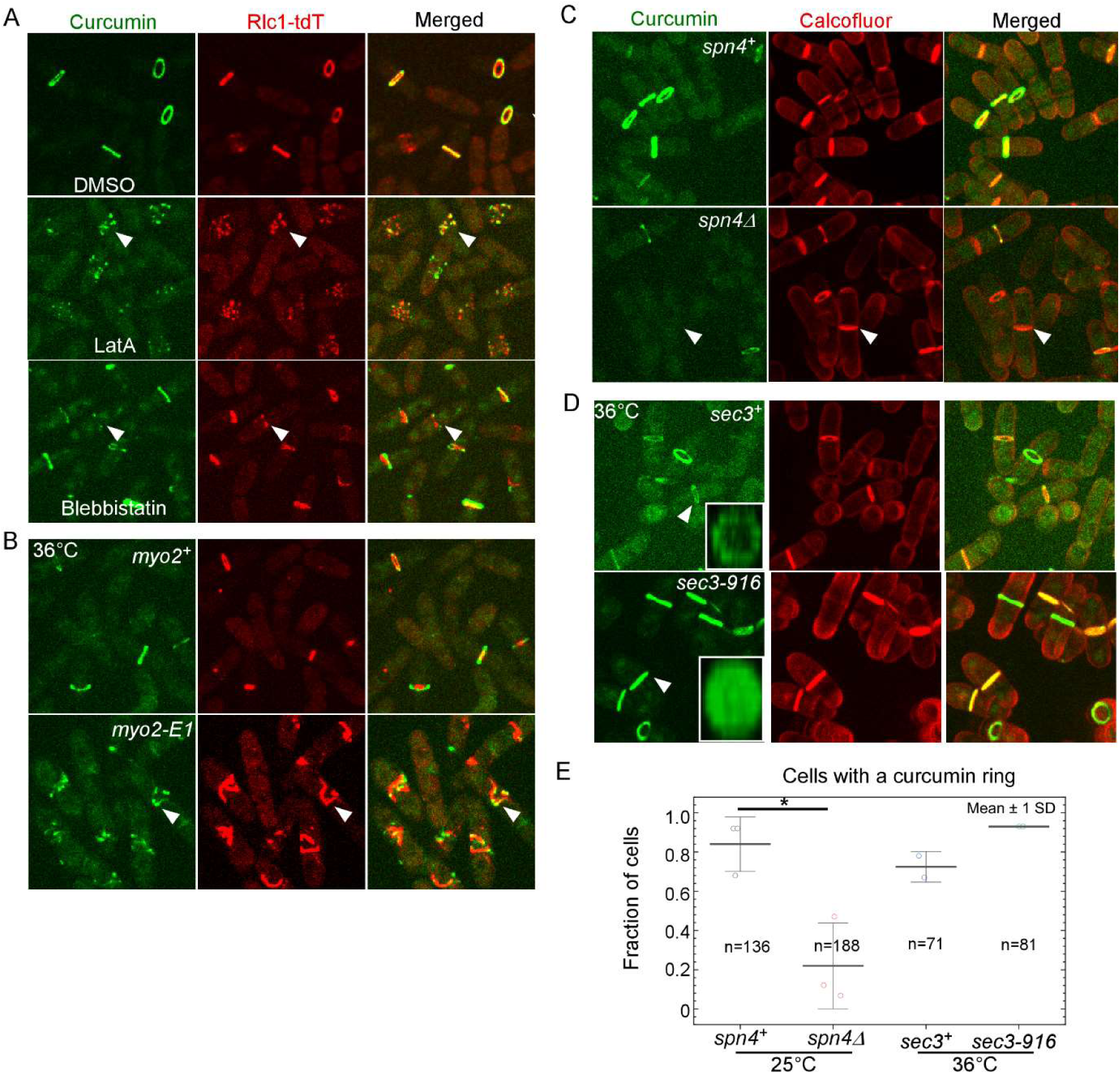
Formation of the curcumin ring depends on the septin ring. 2µM of curcumin (green) was present in all the samples. (A-B) The actomyosin contractile ring is dispensable for curcumin to target the equatorial division plane. (A) Micrographs of the cells expressing Rlc1-tdT (red) in the presence of either DMSO (top), or 10µM latrunculin A (middle), or 20µM blebbistatin (bottom). (B) Micrographs of the wild type and *myo2-E1* cells expressing Rlc1-tdT (red) at the restrictive temperature of 36°C. Arrowhead: the cytoplasmic puncta that was the remnant of the disassembled contractile ring. (C-D) The septins are important for the targeting of the equatorial division plane by curcumin. (C) Micrographs of the wild type (*spn4^+^*) and *spn4Δ* in the presence of calcofluor (red, to stain the septum) Arrowhead: a dividing *spn4Δ* cell was without the curcumin ring. (D) Micrograph of the wild type (*sec3^+^*) and *sec3-916* cells in the presence of calcofluor at the restrictive temperature of 36°C. Inserts: 3D reconstructed head-one of curcumin rings (arrowhead). (E) Dot plots comparing the fraction of the curcumin ring positive cells among the yeast strains. *: P<0.05. Statistical analysis was carried out using Two-tailed unpaired Student’s *t* tests. Representative micrographs from three independent biological repeats are shown. Number of analyzed cells are provided for each sample. Bar: 5 µM.

We next determined whether the septins are required for the formation of the curcumin ring at the equatorial plane, motivated by the strong co-localization between the septin ring and curcumin. In the deletion mutant of *spn4* one of the septin genes that is essential for the septin ring, curcumin disappeared from the equatorial plane of most dividing cells (Fig. 3C). Among the dividing *spn4Δ* cells, marked by calcofluor, the curcumin ring was found in only 22% of them, far less than the wild type cells (84%) (Fig. 3C and E). Even the few remaining curcumin rings appeared to be less intense in the septin mutant cells than they were in the wild type cells. Next, we looked into the localization of curcumin in the exocyst mutant. Since the exocyst regulates the localization of the septin ring (Singh et al., 2025), we hypothesized that the localization of curcumin shall be altered in the exocyst mutant cells as well. In the temperature-sensitive exocyst mutant *sec3-916*, the septin ring becomes more intense and distributes more broadly at the division plane than it does in the wild type cells (Singh et al., 2025). At the restrictive temperature of 36°C, in the *sec3-916* cells, the curcumin ring was still found in most of the dividing cells (93%), actually higher than the wild type cells (72 %) (Fig. 3D and E). However, curcumin, instead of encircling the division plane as a ring, spread out on the division plane evenly as a disc in these mutant cells (Fig. 3D). Not surprisingly, the fluorescence intensity of curcumin at the division plane increased seven times in these exocyst mutant cells (4,423±1,590, average±standard deviations, n=29), compared to those in the wild type cells (633±400, n=31) (Fig. 3D). Therefore, we concluded that the septin cytoskeleton is required for curcumin to both target the division plane and to form a ring there.

The targeting of the equatorial division plane by curcumin raised the question of whether this small molecule interferes with fission yeast cytokinesis. To answer this question, we first measured the assembly, maturation and constriction of the contractile ring in the presence of curcumin, based on time-lapse fluorescence microscopy of the dividing cells (Fig. 4A). Without the presence of curcumin, the yeast cells took ∼20 mins to assemble a complete contractile ring, marked by Rlc1-tdTomato, from the cytokinetic nodes (Fig. 4B), consistent with the previous studies (Wu et al., 2006). Adding curcumin, ranging from 2µM to 8 µM, did not significantly alter the duration of this step of cytokinesis (Fig. 4B). The following step, the ring maturation, from the completion of the ring assembly to the start of the ring constriction, took an average of 12 mins without curcumin (Fig. 4C). Again, this was not significantly altered by the presence of up to 8μM of curcumin (Fig. 4C). In contrast to these first two steps, the rings constricted significantly more slowly (-20%, P<0.001) at a rate of 0.28±0.03 µm/min (average ± S.D.) even in 2µM curcumin than those did so without curcumin (0.35±0.04 µm/min) (Fig. 4D). Increasing the curcumin concentration to 8µM reduced the rate further (-49%) to 0.18±0.07 µm/min (Fig. 4D). We concluded that low micromolar concentration of curcumin inhibits the cleavage furrow ingression during cytokinesis.

**Figure 4.**
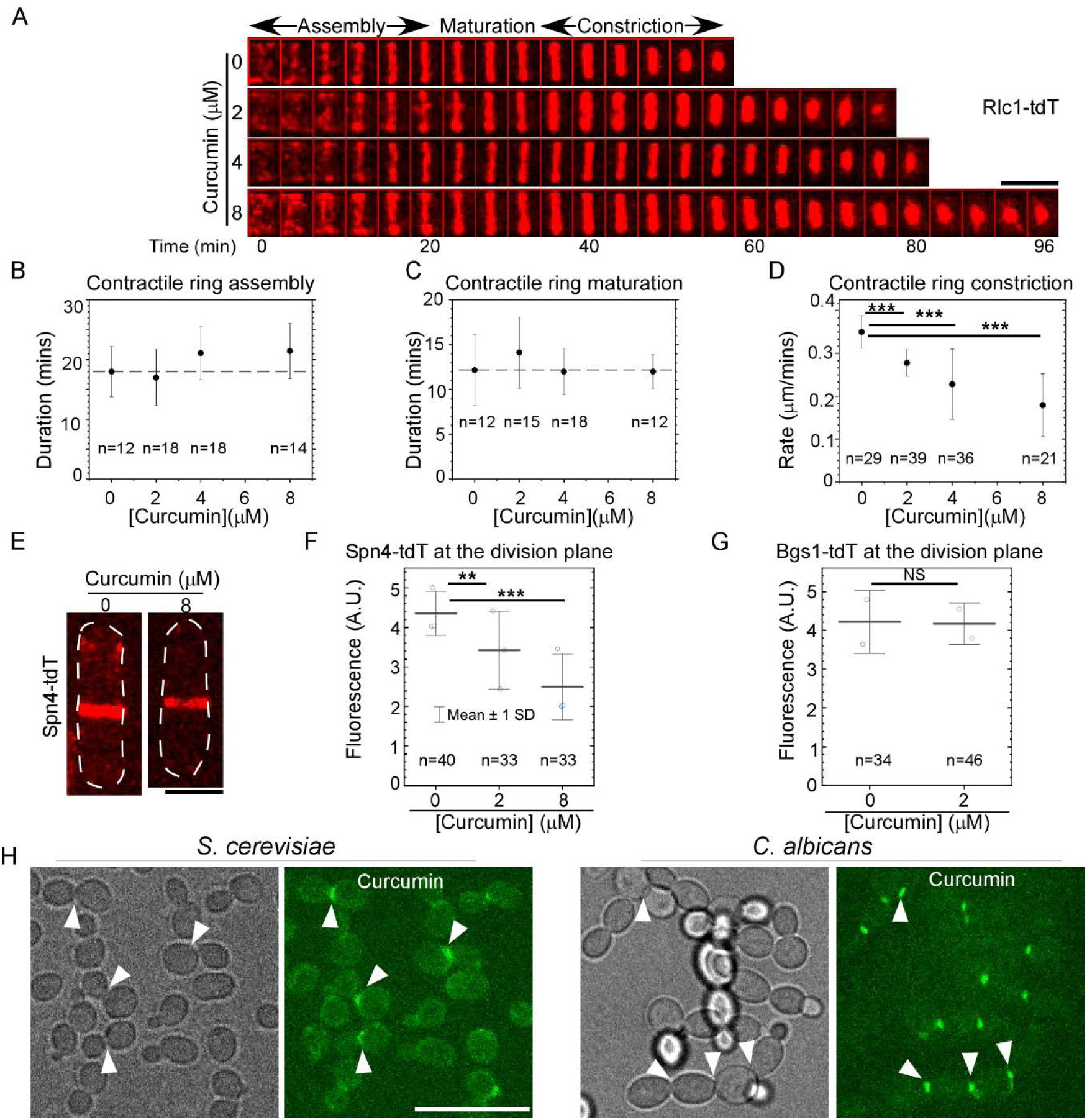
High concentrations of curcumin slow down the contractile ring constriction and curcumin similarly marks the division plane of two other yeasts, *S. cerevisiae* and *C. albicans*. (A-D) Dosage-dependent effect of curcumin on fission yeast cytokinesis. (A) Time-series of the division plane of the cells expressing Rlc1-Td (red) in the presence of 0-8 µM of curcumin. Number: time in minutes from the start of contractile ring assembly. (B-D) Scatter plots of the duration of contractile ring assembly (B), maturation (C) and the rate of contractile ring constriction (D) against the curcumin concentrations. The ring assembly and maturation were not slowed down significantly by curcumin (P>0.05), but the ring constriction was. (E-G) The effect of curcumin on the localization of Spn4 and Bgs1 at the division plane. (E) Micrograph of a cell expressing Spn4-tdT (Red) in either DMSO or 8µM curcumin, 2 min before the daughter cell separation. (F-G) Scatter plots of the peak fluorescence intensities of Spn4-tdT (F) and Bgs1-tdT (G) at the division plane respectively. (H) Micrographs of the wild-type *S. cerevisiae* (left) or *C. albicans* (right) cells in the presence of 2µM curcumin (green) after 10-minute incubation. Arrowhead: the curcumin band. Representative micrographs from three independent biological repeats are shown. *: P<0.05. **: P<0.01***: P<0.001, NS: not significant. Statistical analysis was carried out using Two-tailed unpaired Student’s t tests. Number of the analyzed cells are provided for each sample. Bar: 5 µM.

Next, we explored the potential mechanism by which curcumin inhibits cytokinesis by examining the localization of septins through time-lapse microscopy of the cells expressing Spn4-tdTomato. The peak fluorescence intensity of Spn4-tdTomato at the equatorial division plane reduced by 43% (P<0.001) in the presence of 8µM curcumin (Fig. 4E-F). Even 2µM curcumin reduced the fluorescence by 21% (P<0.01) (Fig. 4F). In contrast to the septins, the peak fluorescence intensity of the glucan synthase Bgs1-tdTomato at the cleavage furrow did not change significantly in the presence of 2µM curcumin (Fig. 4G). Therefore, curcumin hinders the localization of septins at the division plane.

Lastly, we determined whether curcumin similarly targets the division plane of other yeasts cells. We picked two other commonly used model organisms, the Baker’s yeast *S. cerevisiae* and the pathogenic yeast *C. albicans*. As in the fission yeast cells, curcumin marked the bud neck of dividing *S. cerevisiae* cells at 2µM (Fig. 4H). However, compared to those in fission yeast, these curcumin bands in budding yeast cells were less intense. A much brighter curcumin band decorated the bud neck of dividing *C. albicans* cells (Fig. 4H). We concluded that curcumin targets the division plane of two other yeasts as well, suggesting that the cytokinetic curcumin ring may be a shared feature of yeasts.

It is surprising that our study revealed for the first time the existence of an equatorial curcumin ring in yeast cells, considering the large number of studies that have examined the effect of curcumin towards fungi including yeasts. This could be due to two reasons. First, the past studies typically examined the effects of curcumin at concentrations much higher, often exceeding 100µM (Minear et al., 2011), than ours (2µM). At such high concentration, curcumin, with a molecular weight of only 368 Da, may have bound many other intracellular structures nonspecifically. Consistent with the prediction, we also observed that curcumin stained the cytoplasm of some cells in addition to the equatorial ring at an elevated concentration of 8µM. The other critical factor that may have worked to our favor is timing. We observed the yeast cells just a few minutes after labeling them with curcumin. In comparison, past studies typically waited for hours after addition of curcumin. This again may have allowed curcumin to bind either other non-specific or low-affinity intracellular targets. Overall, non-specific binding of other intracellular targets by curcumin when its concentration is high may have prevented the observation of the curcumin band in those earlier studies.

Our work strongly suggests that the yeast septin ring is one of the main targets of curcumin. Fission yeast cells possess seven septins, four of which Spn1, 2, 3 and 4 express in vegetative cells and oligomerize into filaments (Longtine et al., 1996; Onishi et al., 2010). During cell division, these septins form a ring at the division plane (Singh et al., 2025; Tasto et al., 2003) to promote the daughter cell separation (An et al., 2004). We found several lines of evidence supporting that the yeast septin cytoskeleton is a main target of curcumin. First, the septin ring co-localized with the curcumin ring strongly at the equatorial division plane. Secondly, both curcumin and septin rings appeared at the division plane at roughly the same time, just before the contractile ring starts to constrict. Both maintained their diameter until the end of cytokinesis when they split into two one in each daughter cell. No other component of the fission yeast cytokinetic machinery behaves as such. Thirdly, deletion of one of the septins *spn4*, reduced the localization of curcumin to the cleavage furrow significantly. And in the exocyst *sec3* mutant, the altered distribution of the curcumin ring at the division plane mirrors that of the septin ring described in an earlier study (Singh et al., 2025). Fourthly, curcumin hinders the localization of Spn4 at the equatorial division plane. In addition to our findings, a genome-wide screen of in *S. pombe* mutants found that the deletion mutant of another septin *spn3* is hyper-sensitive to a derivative form of curcumin hydrazinocurcumin (Baek et al., 2008). Overall, our data points to the yeast septin ring as a strong target of curcumin at the equatorial division plane, suggesting that curcumin can be used as a potential fluorescent probe for the septin cytoskeleton in yeasts.

Targeting the yeast septin structure with curcumin can hold an advantage for the plant that synthesizes this small molecule. Septin is ubiquitous in fungi, animals and algae, but it is completely absent from the land plants (Kinoshita, 2003). By synthesizing curcumin in their roots, *C. longa* may gain the advantage of targeting a cellular structure unique to fungi, which the plant has to compete against in the soil. Our findings may explain the value of curcumin to C. *longa*, as a defense mechanism targeting fungal cells specifically.

In addition to septins, curcumin may also target the membrane recognized by the septin cytoskeleton. Even in the deletion mutant of *spn4* which is essential for the septin ring assembly (An et al., 2004)*, ∼*20% of dividing cells still possessed the curcumin ring, albeit one that was substantially less intense than those in the wild type cells. This suggests that curcumin may recognize a minor target at the division plane. Septins recognize curved membrane through their direct interaction with the lipids in the plasma membrane (Bridges et al., 2016). Targeting of this membrane may explain the result from a previous genetic screen of budding yeast mutants that are hyper-sensitive to curcumin (Minear et al., 2011). It found that lipid biosynthesis, including that of ergosterol and phospholipids, is essential to the resistance of yeast to high concentrations (100µM) of curcumin (Minear et al., 2011). Our hypothesis is also consistent with the lipophilic nature of curcumin (Tonnesen et al., 2002). Overall, it is likely that curcumin recognizes the septin cytoskeleton as the primary target and the membrane as the secondary target at the equatorial division plane.

Curcumin may possess a general inhibitory effect against cytokinesis of yeasts, besides *S. pombe*. This is consistent with the localization of curcumin band at the equatorial division plane of three evolutionary diverse yeast species. The small molecule specifically inhibits the cleavage furrow ingression of fission yeast, by disrupting the septin cytoskeleton organization at the equatorial division plane. This inhibitory effect of curcumin against cytokinesis is consistent with the recent finding that curcumin interferes with cytokinesis of the green algae *Chlamydomonas reinhardtii* (Clark-Cotton et al., 2025). Overall, our work strongly suggests that curcumin delays fission yeast cytokinesis through inhibiting the septin ring.

Taken together, our study revealed the novel capability of curcumin to target dividing yeast cells by forming a ring even at a relatively low concentration. Curcumin targets the septin ring at the equatorial division plane. Formation of the curcumin ring depends primarily on the septins. In the future, curcumin can be exploited to explore for its antifungal potential, considering that this turmeric is well-tolerated in human bodies. For cell biologists, curcumin may also be a valuable fluorescent marker for the septin cytoskeleton at low concentrations.

## Materials and Methods

### Yeast Strains and growth conditions

We followed the standard procedures for yeast cell culture (Moreno et al., 1991). The *S. pombe* yeast cells were inoculated in liquid YE5s media at 25°C in an orbital shaker (Eppendorf, USA). Both *S. cerevisiae* cells and *C. albicans* cells were inoculated in YPD media. The temperature-sensitive mutants were inoculated at 36°C for 4 hours before microscopy.

### Chemicals

Curcumin is commercially available from Millipore Sigma (C1386). A 10mM stock solution was prepared in DMSO and stored at -20°C for long-term usage. The curcumin supplemented yeast media was freshly prepared on the day of experiment by diluting the stock solution in YE5s to the final working concentration of between 2-8µM, through a series of dilutions. Calcofluor White (1mg/ml) was commercially available from Millipore Sigma (18909). Lectin (Sigma, L2380) was prepared in water for 1mg/ml stock solution. Latrunculin A stock solution (1mM) was prepared by adding 240µl of DMSO to 100µg of LatA (Sigma, 428021) and stored at -20°C. Blebbistatin (Sigma, B0560) was prepared as a 2mM stock solution in DMSO stored at -20°C.

### Fluorescence microscopy

For live microscopy throughout this study, we imaged the yeast cells in liquid media in a glass-bottomed 35mm petri dish using a previously described method (Poddar et al., 2021). Briefly, to prepare the dish (Cellvis, USA), we added 50µl of 50µg/ml of lectin to the center of the coverslip and allowed it to dry out slowly at 30°C for 3-4 hours. We then deposited 20µl of the exponentially growing yeast culture to the cover slip and left it to stand for 10 mins. Afterwards, we added 2 ml of YE5s media, supplemented with either 2µM curcumin or 0.02% DMSO (control), to the petri dish and inoculated for another 10 mins before proceeding to microscopy.

We used a spinning disk confocal system controlled by iQ3 software (Andor) for all the microscopy work. It consists of an Olympus IX71 microscope body and a CSU-X1 spinning disk unit (Yokogawa, Japan). It is equipped with a motorized stage and a Piezo Z Top plate (ASI, USA). Images were captured on an EMCCD camera (IXON-897, Andor). The microscope employed solid-state lasers with wavelengths of 488, 561 and 405nm. The fluorescence of curcumin was excited at 488nm, and its emission was captured at 525nm. Unless otherwise noted, a 60x objective lens (Olympus, Plan Apochromat, NA = 1.40) was used. Typically, a Z-stack of 8 slices at an interval of 1µm was acquired. For the temperature-sensitive mutants, we imaged the cells in a temperature-controlled chamber (Okolab, Italy) maintained at 36°C.

### Image analysis

All the image analysis were carried out using NIH ImageJ, its open-sources plugins and our own custom-made macros. We used the ImageJ 3D plugin to reconstruct the head-on view of the cleavage furrow, based on the Z-stack of a dividing cell. The figures were prepared using Canvas X (ACD Systems, USA). The plots were prepared using Origin (OriginLab, USA).

### Counting the fraction of curcumin band positive cells

To count the dividing cells possessing a cytokinetic curcumin ring, the fission yeast cells were incubated in the YE5s media supplemented with both curcumin (2µM) and calcofluor (1µM) at the room temperature in a 35mm petri dish with a coverslip bottom for 10 mins. Following the microscopy, we projected a Z-stack into a maximum intensity projection for analysis. We first counted the number of the cells with a calcofluor stained septa using ImageJ Cell Counter. Among these septated cells, we counted those with a curcumin ring at the equatorial division plane. We then calculated the fraction of curcumin ring-positive cells based upon these two measurements.

### Quantifying the curcumin fluorescence at the division plane

We first projected the Z-stacks into average intensity projections. To measure the localization of curcumin at the division plane, we quantified the average fluorescence in 0.8µm wide by 3.8µm long zone at the equatorial plane of a dividing cell. All the fluorescence intensity measurements were subtracted by the camera noise, quantified as the fluorescence intensity in a field where no cells were present.

### Quantitative analysis of cytokinesis

To measure cytokinesis in the presence of curcumin, we first added the YE5s media supplemented with either DMSO (0.02%) or various concentrations of curcumin to the cells cultured in a 35mm glass-bottomed petri dish. The cells were then incubated in the dark for 30 minutes, followed by live microscopy. We typically took two-hour time-lapse series with two-minute intervals at the room temperature (∼23°C).

For analysis, the Z-stacks were first projected as maximum intensity projections. The duration of the contractile ring assembly was counted from when the cytokinetic nodes appeared at the division plane to when a full ring materialized at the division plane. The duration of the ring maturation was counted from when the full ring was assembled to when the ring started to constrict. To quantify the rate of contractile ring constriction, we first constructed the fluorescence kymograph for each constricting ring based upon the time-series. The rate was calculated as the circumference of the ring divided by the duration of the ring constriction. The circumference of a dividing cell was calculated based on the diameter of the ring measured at the start of its constriction. The duration of the ring constriction was counted from when the ring started to narrow till when the ring constricted completely with its remnant diffusing away.

### Quantifying the localization of Spn4 and Bgs1 at the division plane

We first collected time-lapse series of the cells expressing either Spn4-tdTomato or Bgs1-tdTomato in the presence of either curcumin or DMSO at the room temperature for two hours with an interval of 2mins. We projected the Z-stacks into average intensity projections. To measure the localization of either Spn4-tdTomato or Bgs1-tdTomato at the division plane, we quantified their respective average fluorescence in a 0.8µm wide zone along the equatorial plane of a dividing cell. For measuring the peak fluorescence intensity of Spn4-tdTomato at the division plane, we took the measurement two minutes prior to the daughter cell separation, because the fluorescence intensity of Spn4-tdTomato remained roughly unchanged for about 20 mins before the cell separation. To measure the peak fluorescence intensity of Bgs1-tdTomato, we took the measurement at the division plane, 28 mins before the daughter cell separation. The fluorescence intensity measurements were subtracted by the camera noise, quantified as the fluorescence intensity in a field where no cells were present.

### Pharmacological treatment

To disassemble the actomyosin contractile ring, 5µl of either LatA (1mM) or blebbistatin (2mM) stock solution was added to 495µl of exponentially growing yeast cells in a microcentrifuge tube. The cells were inoculated in the dark at room temperature for 1 hour. For the control, DMSO was added in the place of the drugs. Afterwards, 10µl of curcumin (100µM) was added to the culture, followed by 20 mins of inoculation in the dark. 20µl of the treated cell culture was deposited onto a glass-bottomed 35mm petri dish. The cells were allowed to attach for 10 mins in the dark before microscopy.

**Table 1:**
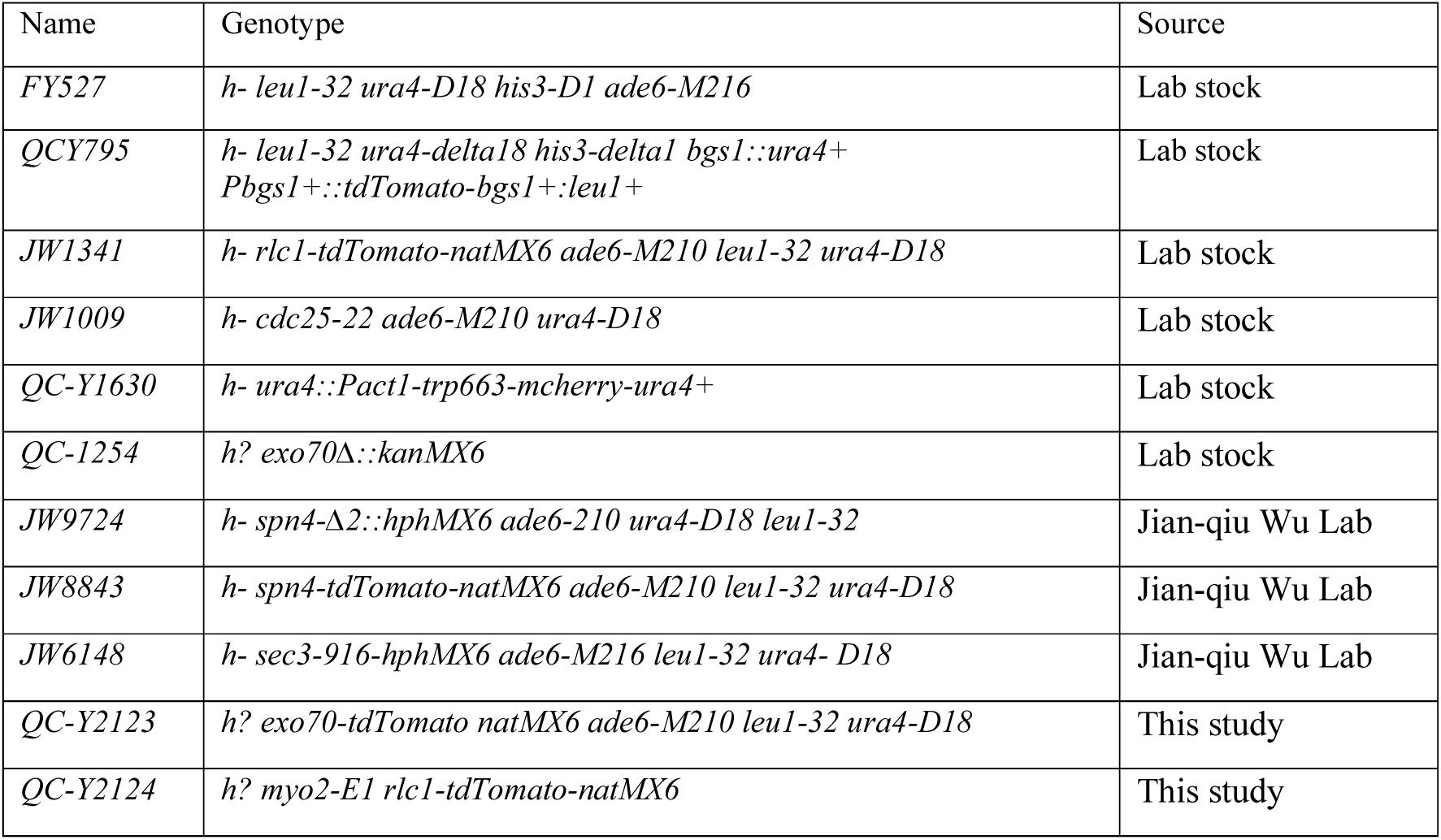
List of fission yeast strains used in this study.

## Acknowledgements

The authors thank the Chen lab members for their technical support. We thank Tatiana De Souza from the Heather Conti lab (The University of Toledo) for sharing the wild-type *C. albicans* culture and Tony Hazbun (Purdue University) for sharing the wild-type *S. cerevisiae* strain. We thank Jian-qiu Wu (Ohio State University) and Kathy Gould (Vanderbilt University) for kindly sharing their fission yeast strains.

## Author Contributions

Varmila Kulasegaram: *Investigation, Formal analysis, Methodology, Writing - review & editing*

Daphne Alves Dias: *Investigation, Formal analysis, Funding acquisition, Methodology, Writing - original draft, Writing - review & editing*

Anna Okorokova-Façanha: *Funding acquisition, Writing - original draft*

Qian Chen: *Investigation, Formal analysis, Conceptualization, Funding acquisition, Project administration, Writing - original draft, Writing - review & editing*

